# Metal transporter SLC39A14/ZIP14 modulates regulation between the gut microbiome and host metabolism

**DOI:** 10.1101/2021.12.22.473859

**Authors:** Trista L. Thorn, Samuel B. Mitchell, Yongeun Kim, Min-Ting Lee, Janine M. C. Comrie, Elizabeth L. Johnson, Tolunay B. Aydemir

## Abstract

Zinc (Zn) plays a critical role in maintaining intestinal homeostasis by regulating intestinal epithelial cells, host immune cells, and gut microbiome community composition. Deletion of metal transporter *Slc39a14/Zip14* causes spontaneous intestinal permeability with low-grade chronic inflammation, mild hyperinsulinemia, and greater body fat with insulin resistance in adipose, suggesting a role for ZIP14-mediated intestinal metal transport in regulating both intestinal homeostasis and systemic metabolism. Here, we showed the function of ZIP14-mediated Zn transport in the gut microbiome composition and how ZIP14-linked changes to gut microbiome community composition are correlated with changes in host metabolism. Deletion of *Zip14* generated Zn-deficient epithelial cells and luminal content in the entire intestinal tract; reduced bacterial diversity and *Saccharomyces cerevisiae* (*S. cerevisiae*) overgrowth; altered host metabolome; and shifted host energy metabolism toward glucose utilization. This work provides evidence for the regulation of gut microbiome composition, host metabolome, and energy metabolism by metal transporter ZIP14.

**Significance:** Intestinal permeability, gut dysbiosis, and Zn dyshomeostasis are emerging signatures of inflammatory bowel diseases and metabolic disorders such as type-2-diabetes and obesity. Zn deficiency is a common clinical finding among these diseases. Zn is essential for the regulation of the intestinal epithelial cells, host immune cells, and the gut microbiome. Transporter-mediated mobilization of Zn plays a critical role in maintaining intestinal homeostasis by facilitating the targeted tissue/cell-specific function. However, studies are lacking in linking transporter-mediated Zn mobilization, gut microbiome, host’s intestinal health, and metabolism. Using the systems-level approach, this study revealed novel findings that deletion of *Slc39a14/Zip14* resulted in altered intestinal Zn homeostasis, gut microbiome composition, host metabolome and energy metabolism.

## Introduction

The essential trace metal Zn is the second most abundant metal in the body, with diverse and indispensable catalytic, structural, and regulatory functions ^1^. Zn transporters maintain Zn homeostasis by facilitating the cellular and subcellular distribution of Zn to and within tissues. Two families of metal transporters, SLC39A/ZIP1-14 (Zrt-Irt-like proteins) and SLC30A/ZnT1-10 (Zn transporter), have opposing functions. ZIP family transporters are responsible for the transport of metals into the cytoplasm from the extracellular space and within the lumen of intracellular compartments. ZnT family transporters transport metals from the cytoplasm to either extracellular space or intracellular compartments. Intestinal Zn transport is central to both intestinal and systemic Zn homeostasis. Zn is absorbed mainly by the function of apically localized Zn transporter, ZIP4 ^2–4^. On the basolateral side, ZnT1 and, ZIP5 and ZIP14 are proposed to transport Zn from the enterocyte into the circulation and from circulation to enterocytes, respectively ^5,6^. Metallothioneins and the ZnT2, ZnT5A, ZnT6, ZnT7, ZIP7, and ZIP14 transporters provide intracellular Zn movement ^6–9^.

Emerging evidence suggests that Zn transporter-mediated mobilization of Zn plays a critical role in maintaining intestinal homeostasis by facilitating the targeted tissue/cell-specific function. Zn is essential for the regulation of the intestinal epithelial cells, host immune cells, and intestinal commensal bacteria ^10^. Mice lacking ZIP4 in the intestinal epithelial cells showed loss of Zn in Paneth cells with impaired proliferation and epithelial architecture ^11^. Deleting ZIP7 in the intestinal epithelial cells in mice caused defects in stem cell survival, the proliferation of crypts, and resolving endoplasmic reticulum stress ^12^. ZnT2 is expressed on the secretory granules in Paneth cells, promotes the accumulation of zinc, and regulates the secretion of antimicrobial molecules ^13^. Genome-wide association studies revealed an association between a variant of Zn transporter ZIP8 and type of inflammatory bowel disease (IBD), Crohn’s disease (CD) with altered gut microflora ^14^. Intestinal permeability and inflammation are signatures of IBD. In a humanized mouse model of the *ZIP8* variant, enhanced disease susceptibility was reported in chemically induced colitis ^15,16^.

We have previously shown that deletion of *Zip14*, the phylogenetically closest relative of *Zip8* ^17^, induced spontaneous intestinal permeability with low-grade chronic inflammation (metabolic endotoxemia), enlarged islets with mild compensatory hyperinsulinemia, and greater body fat with insulin resistance in adipose tissue ^6,18–20^. Besides Zn, ZIP14 also transports manganese and iron ^21^. We have previously shown that intestinal manganese elimination was impaired in *Zip14* KO mice ^22,23^. Here we present our novel findings on the function of ZIP14-mediated Zn transport in intestinal tissue, microbial profile, and how ZIP14-linked changes in gut microbiome and metabolites modulate host metabolism.

## Results

### Ablation of *Zip14* alters gastrointestinal Zn homeostasis

We first examine the effect of the *Zip14* deletion on Zn homeostasis in the entire intestinal tract using microwave plasma atomic emission spectrometry (MP-AES). In the *Zip14* KO mice, Zn concentrations were lower in the proximal and distal intestine and the colon tissue (Fig. 1A). The total intestinal content is composed of both unabsorbed dietary Zn and Zn from endogenous sources, including transporter-mediated transepithelial flux of Zn in the serosal to the mucosal direction. We collected total intestinal content and separated it into two phases by centrifugation. Luminal content (supernatant) represents ionic and protein-bound Zn, and digesta (pellet) represents the unavailable Zn for host absorption ^24^. Both luminal content and digesta had decreased concentrations of Zn in the *Zip14* KO (Fig. 1B). These data collectively suggested that in the absence of basolaterally localized ZIP14, Zn transport was impaired in serosal to mucosal direction, resulting in a chronic Zn deficiency in the entire GI tract’s tissue, luminal content, and digesta. Zn absorption occurs throughout the intestinal tract as a function of Zn transporters. Since the highest Zn absorption occurs in the proximal intestine, we measured the abundance of known intestinal Zn transporters from isolated enterocytes. We observed more pronounced changes in the female mice. Compared to WT, the abundance of apical Zn importer ZIP4 and basolateral Zn exporter ZnT1 in *Zip14* KO mice was increased and decreased, respectively (Fig. 1C), supporting the Zn-deficient conditions in the enterocytes of *Zip14* KO mice. However, these changes were not sufficient to compensate for the deficiency caused by the absence of ZIP14.

**Figure 1.**
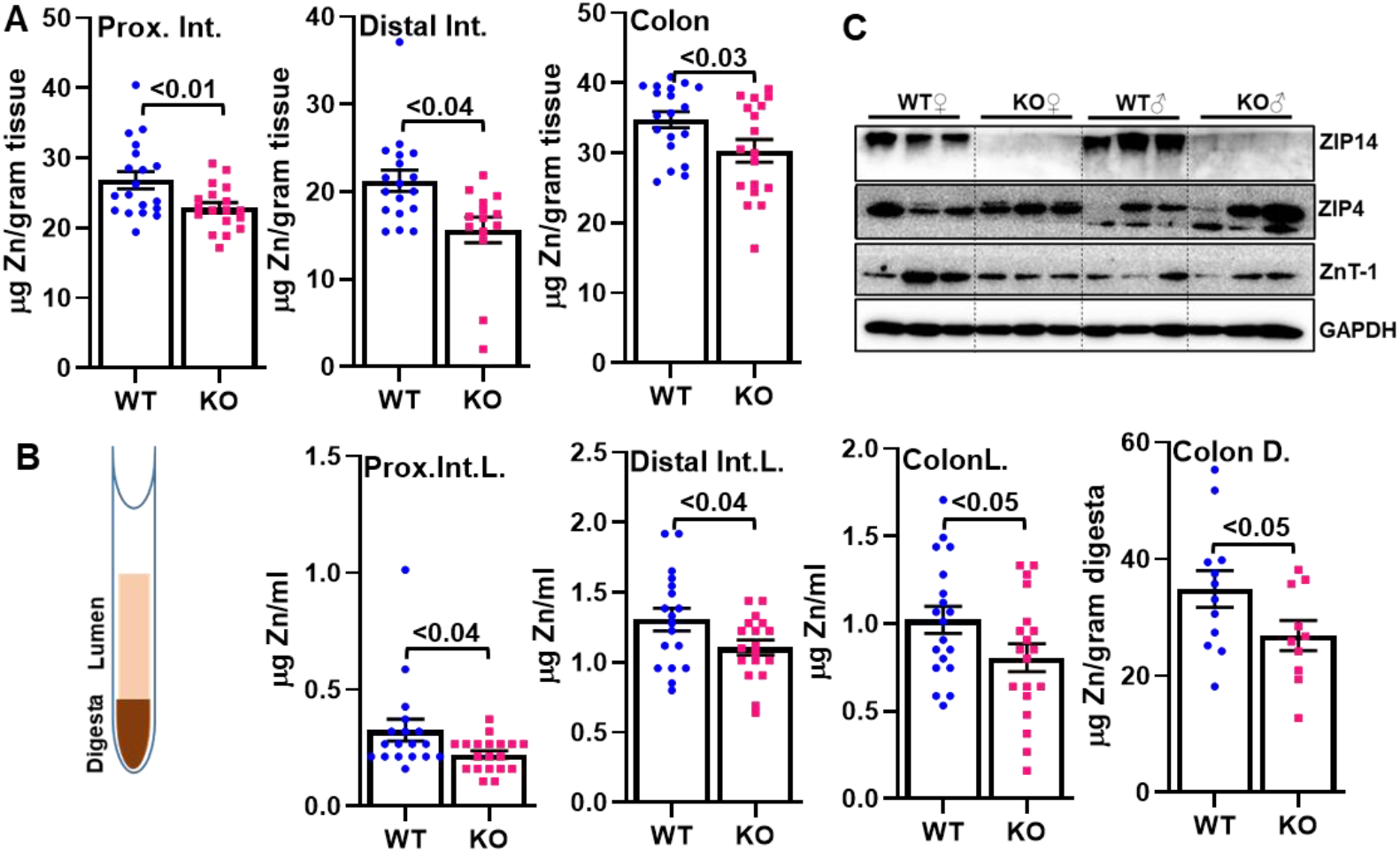
Altered gastrointestinal Zn homeostasis in *Zip14* KO mice. Zn concentrations in gastrointestinal tissues (A) and the luminal contents (B) were measured by MP-AES. Values are means ± SE; n = 16-19 (both female and male mice were included). Student’s t-test for WT vs. WB-KO comparison. C) Abundance of Zn transporters, ZIP14, ZIP4, and ZnT1, in isolated enterocytes was shown by western blot analysis.

### Ablation of *Zip14* modulates gut microbiome, and ZIP14-shaped microbiome can influence intestinal permeability and metabolic phenotypes in the host

Intestinal barrier function and consequent metabolic alterations are regulated by dynamic and reciprocal interactions among the intestinal microbiota, epithelium, and immune system ^25^. Dietary metals are important factors influencing the intestinal microbiome ^26^, as evidenced by the observations that metal availability is crucial for the outcome of host-pathogen interactions. However, much less is known about the effects of dietary metals on commensal bacteria. A limited number of studies have demonstrated that both Zn deficiency and excess can influence the composition of the gut microbiome. However, it is largely unknown whether Zn transporters regulate the gut microbiome through modulating luminal Zn levels. Due to impaired Zn transport (serosal to mucosal), *Zip14* deletion results in lower Zn levels in the gastrointestinal lumen than those in WT mice (Fig. 1B). We hypothesized that a low Zn environment would affect the composition of the gut microbiome. To test this hypothesis, we determined microbiome community composition using 16S rRNA gene sequencing from fecal pellets of *Zip14* KO versus WT mice. We found significantly lower microbiome α-diversity in WB-KO mice when compared to WT mice (Fig. 2A). This suggests that the presence of certain taxa may be reliant on intestinal luminal concentrations of Zn. To better understand which taxa might be susceptible to Zn concentrations in the lumen of the intestine, we looked for differential abundances of taxa between the microbial communities of *Zip14* KO and WT mice (Fig. 2B). We found a decreased abundance of Bacteriodaceae and Clostridiaceae families and increased abundance of Ruminococcaceae families (Fig. 2C), all of which have been observed in IBD patients as well ^27,28^. To better understand how ZIP14-mediated changes in microbiome community composition may function to alter host phenotypes, we transplanted microbiome-depleted mice with fecal contents from *Zip14* KO and WT mice (Fig. 3A). Following the fecal transplant, we have conducted a permeability assay using FITC-Dextran gavage. We found significantly higher levels of FITC-Dextran in the serum of mice that received the fecal slurry from *Zip14* KO mice (Fig. 3B), indicating that the ZIP14-shaped microbiome contributed to the progression of intestinal permeability. Furthermore, the mice that received the fecal slurry from *Zip14* KO were mildly hypoglycemic (Fig. 3C) with significantly greater % body fat (Fig. 3D) and lower % lean mass (Fig. 3E), recapitulating the metabolic phenotype of *Zip14* KO mice ^20^. Collectively these data indicated that the ZIP14-shaped microbiome could influence metabolic phenotypes in the host.

**Figure 2.**
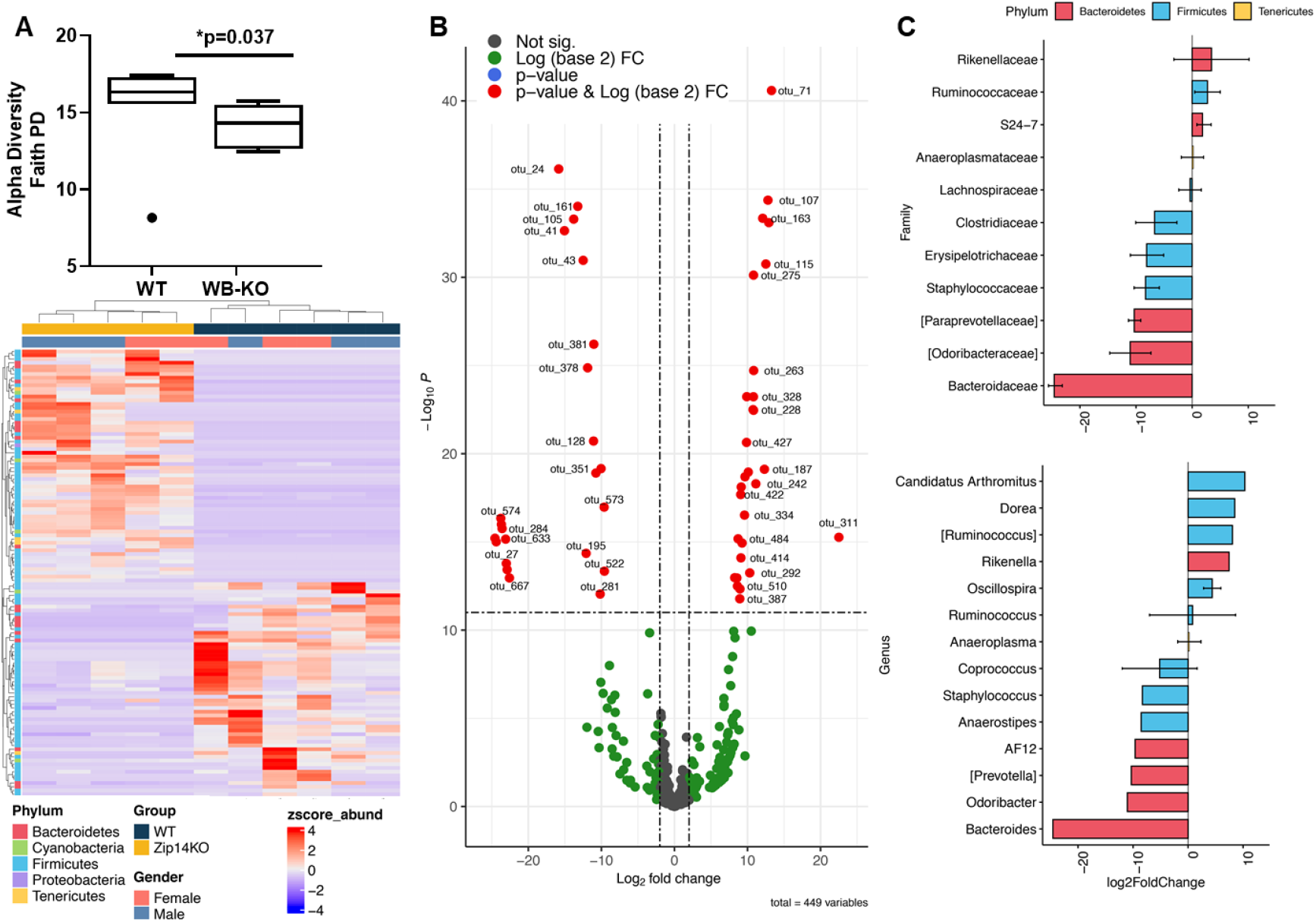
Decreased gut microbial diversity in the WB-Zip14 KO mice. A, B) WB-Zip14 KO displayed decreased α-diversity, as shown by 16S rRNA gene sequencing. Fold change of >2 and p-value < 0.05 were considered significantly different. C) The most significant changes at the family and genus levels (WB-KO/WT) (n=7-8). The details of the analysis are provided in the methods section.

**Figure 3.**
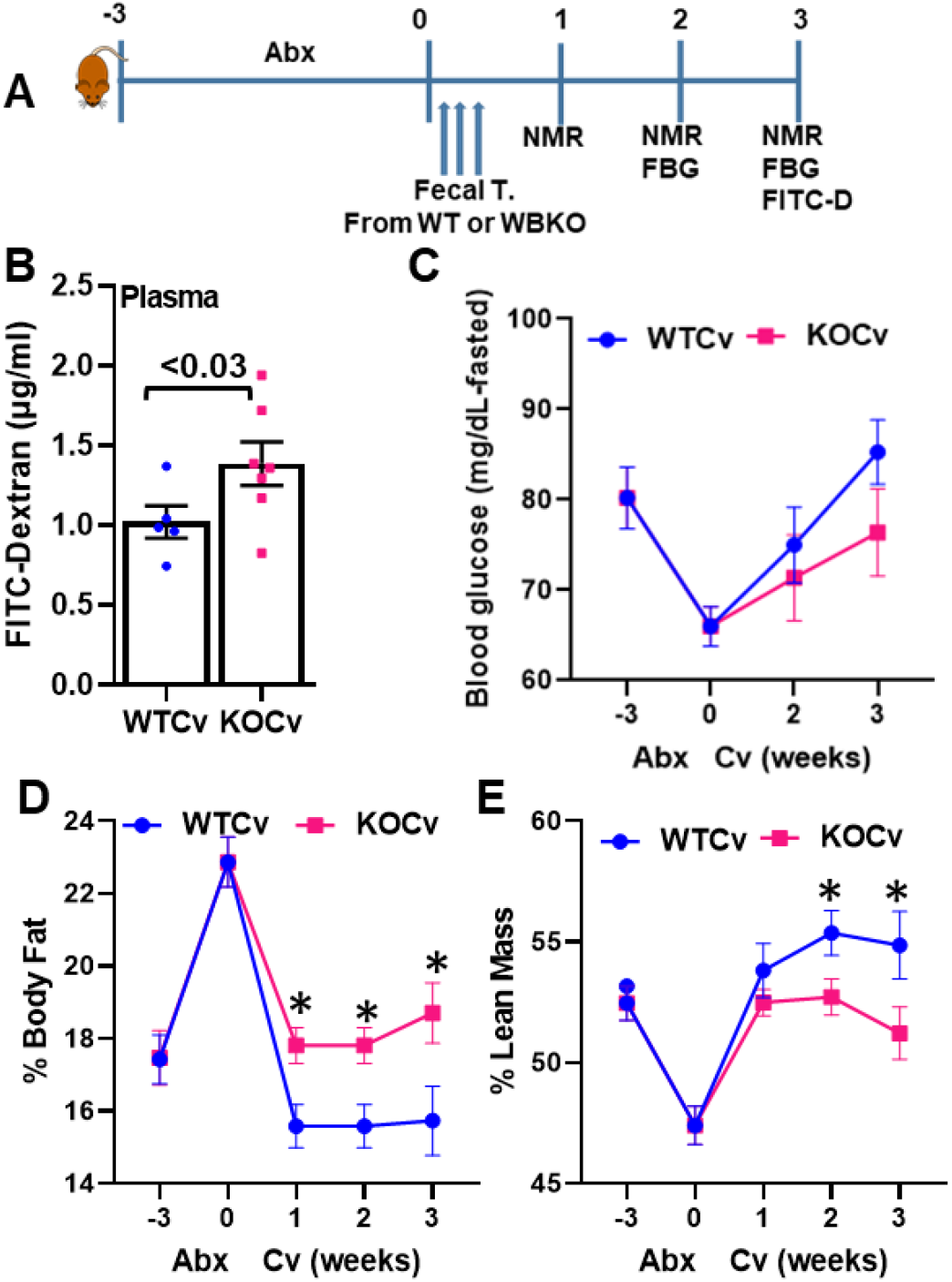
The ZIP14-shaped microbiome can influence intestinal permeability and metabolic phenotypes in the host. A) Following three weeks of antibiotics treatment, mice have received a fecal transplant from either WT (WTCv) or WB-KO (KOCv) mice. B) Plasma FITC-Dextran concentrations were measured fluorometrically. B) Fasting blood glucose was measured by tail bleeding using the OneTouch® Ultra Blood glucose monitoring system. Percent body fat (C) and percent lean mass (D) were assessed by nuclear magnetic resonance. Values are means ± SE; n = 5-7 (both female and male mice were included). Student’s t-test.

### *Saccaramicea cerevisiae (S. cerevisiae)* overgrowth may contribute to intestinal permeability in *Zip14* KO mice

The gut microbiota produces a broad range of metabolic products that impact host biology. To identify the host and microbial-originated metabolites in WT and KO mice serum, we conducted untargeted metabolomics using Gas chromatography (GC) coupled to time-of-flight mass spectrometry (TOF-MS). Principal component analysis (PCA) of overall serum samples showed that metabolite abundance in *Zip14* KO mice significantly differed from WT mice (Fig. 4A), indicating that the *Zip14* KO group could be distinguished from WT according to their different metabolic profiles. Metabolites with a fold change of >1.5 and p-value < 0.05 were considered significantly different (Fig. 4B). The serum concentration of 11 identifiable metabolites varied significantly between WT and WB-KO (Fig. 4B, C). Serum adenosine, serotonin, β-sitosterol, creatinine, N-acetyl-D-tryptophan, and saccharic acid were significantly greater, and only linoleic acid was significantly lower in the serum of *Zip14* KO mice (Fig. 4C).

**Figure 4.**
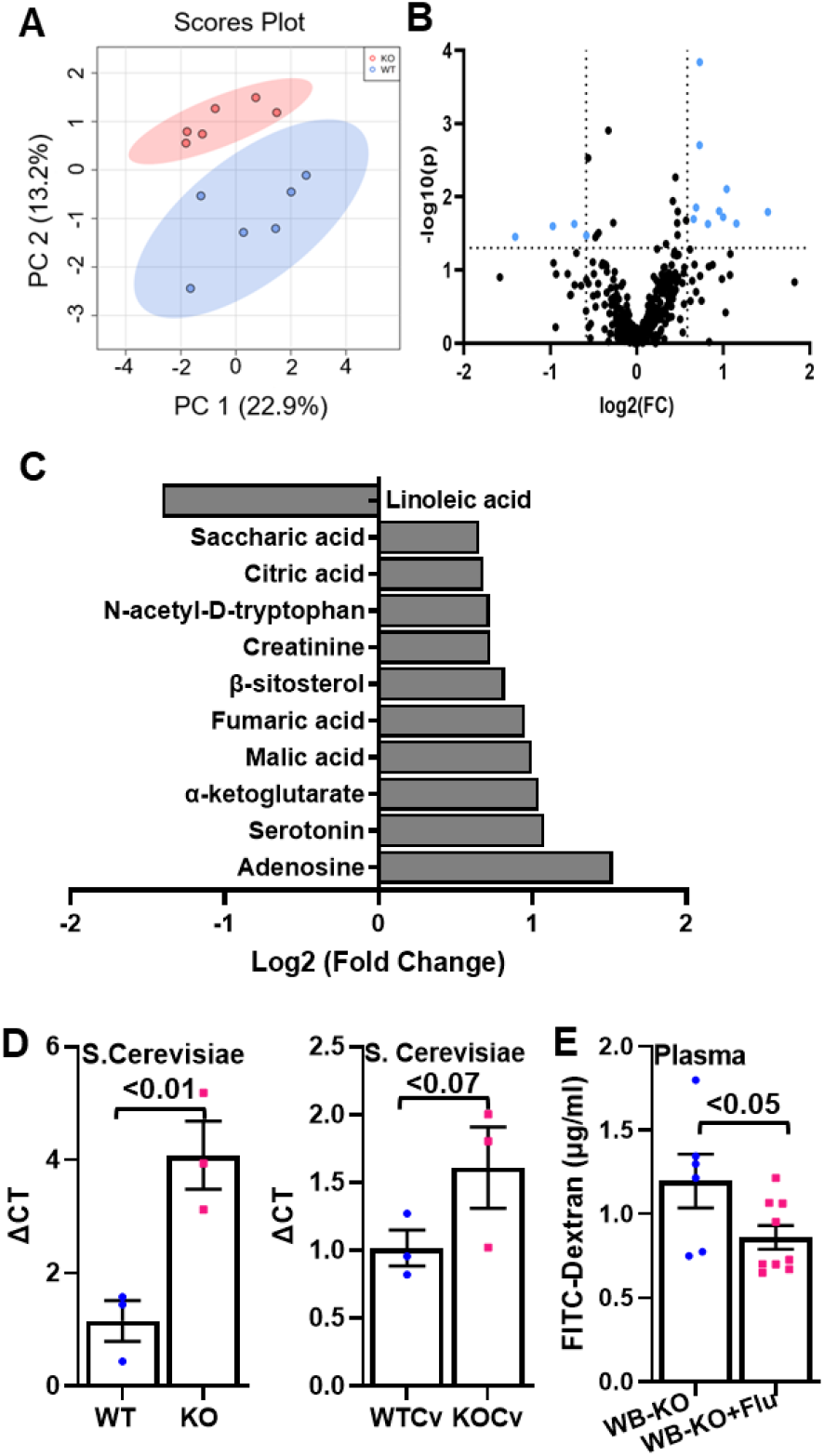
Serum untargeted metabolomics profiling of *Zip14* KO and WT mice. Serum metabolites were measured Gas chromatography (GC) coupled to time-of-flight mass spectrometry (TOF-MS) (n=6 per group). A) The PCA score plot of serum samples from WT and WB-KO mice. B) Volcano plot. C) The serum metabolites with the known identity were statistically different. D) S. Cerevisiae DNA was quantified by qPCR in colonic digesta from either WT and WB-KO or WT mice that received fecal transplant either from WT (WTCv) or KO (KOCv) mice. Values are means ± SE; n = 3. Student’s t-test. E) Following ten days of antifungal (fluconazole) treatment, plasma FITC-Dextran concentrations were measured fluorometrically. Values are means ± SE; n = 6-9. Student’s t-test.

The upregulated metabolite, N-acetyl-D-tryptophan, in the serum of *Zip14* KO mice could be unique to *S. cerevisiae* in the gut due to the production of a unique enzyme acting on D-amino acids, D-amino acid N-acetyltransferase ^29,30^. We hypothesized that the Zn deficient gut in the *Zip14* KO mice with lower bacterial α-diversity could provide the opportunity for yeast overgrowth. Our qPCR results revealed a greater amount of *S. cerevisiae* DNA in the feces of WB-KO mice when compared to their controls (Fig. 4D). To investigate the direct effect of the gut microbiome on the metabolic phenotype of *Zip14* KO mice, we conventionalized antibiotics (abx)-treated WT mice with fecal slurry from either WT (WTCv) or *Zip14* KO (KOCv) mice. We measured *S. cerevisiae* DNA following conventionalization. The KOCv mice had a greater amount of S. cerevisiae; however, it did not reach the statistical significance (p<0.07) (Fig. 4D). To understand the effect of *S. cerevisiae* on the increased intestinal permeability phenotype of WB-KO mice, the ‘mice’s drinking water was supplemented with the antifungal drug fluconazole for ten days. We found a significant reduction in plasma FITC-Dextran levels when WB-KO mice were treated with fluconazole, suggesting that *S. cerevisiae* overgrowth along with reduced bacterial diversity in the WB-KO contributed to the development of intestinal permeability.

### Host energy metabolism is shifted towards continuous utilization of glucose as an energy source in *Zip14* KO mice

To assist in the biological interpretation of our metabolomics data, we conducted pathway and enrichment analyses using MetaboAnalyst ^31^ to explore whether significantly differing metabolites belong to a common pathway. There was an enrichment of significantly differing compounds in several metabolic pathways (Fig. 5A). Among them, the Tricarboxylic Acid Cycle (TCA) cycle pathway had the most significant impact score. TCA cycle intermediate metabolites, citric acid, α-ketoglutaric acid, fumaric acid, and malic acid were significantly upregulated in the serum of *Zip14* KO mice. Of note, a-ketoglutarate, fumaric acid, and malic acid are also responsible for the “alanine, aspartate, and glutamine metabolism ^32^. Disease signature analysis was conducted using metabolite sets reported in human blood. In agreement with the presence of intestinal permeability and low-grade chronic inflammation in *Zip14* KO mice, the top signatures were septic shock and diabetes (Fig. 5B).

**Figure 5.**
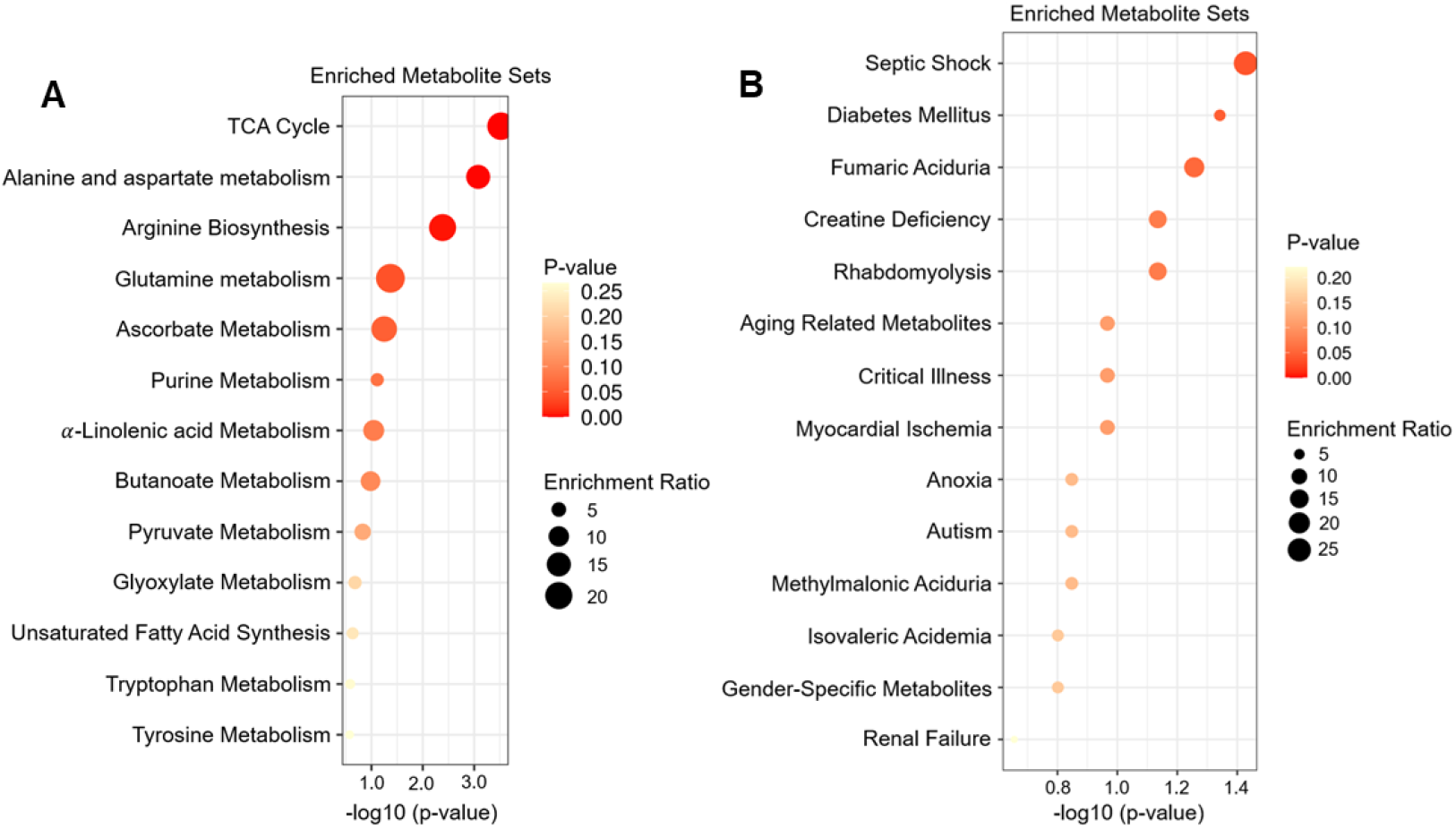
Pathway and enrichment analyses using MetaboAnalyst. For the significantly different metabolites (details in method section), pathway and enrichment analyses were conducted using MetaboAnalyst. A) Pathway analysis was conducted using KEGG human metabolic pathways. B) Disease signature analysis was conducted using metabolite sets reported in human blood.

Despite the increased intestinal permeability, low-grade chronic inflammation, and insulin resistance in adipose, *Zip14* KO mice do not develop hyperglycemia. In addition, our metabolomics data revealed the most significant impact score was on TCA cycle (Fig. 5A), collectively suggesting that *Zip14* deletion may cause a shift in host energy metabolism. To quantify these observed effects on whole-body metabolism, we conducted metabolic phenotyping by using a Comprehensive Laboratory Animal Monitoring System (CLAMS). In the light cycle, oxygen consumption levels (Fig. 6A), carbon dioxide production (Fig. 6B), and respiratory exchange ratio (Fig. 6C) were greater in the *Zip14* KO mice than WT. These data suggested that glucose was preferentially used as an energy source in the *Zip14* KO mice. This may partially explain the prevention of hyperglycemia phenotype. Next, we fed mice with HFD for 16 weeks to force fatty acids (consumption of excess). At the end of the study, fasting blood glucose levels were elevated in HFD fed WT mice, but there was no increase in the HFD-fed *Zip14* KO mice, even though both WT and KO had an equal increase in percent body fat in the HFD fed group (data not shown).

**Figure 6.**
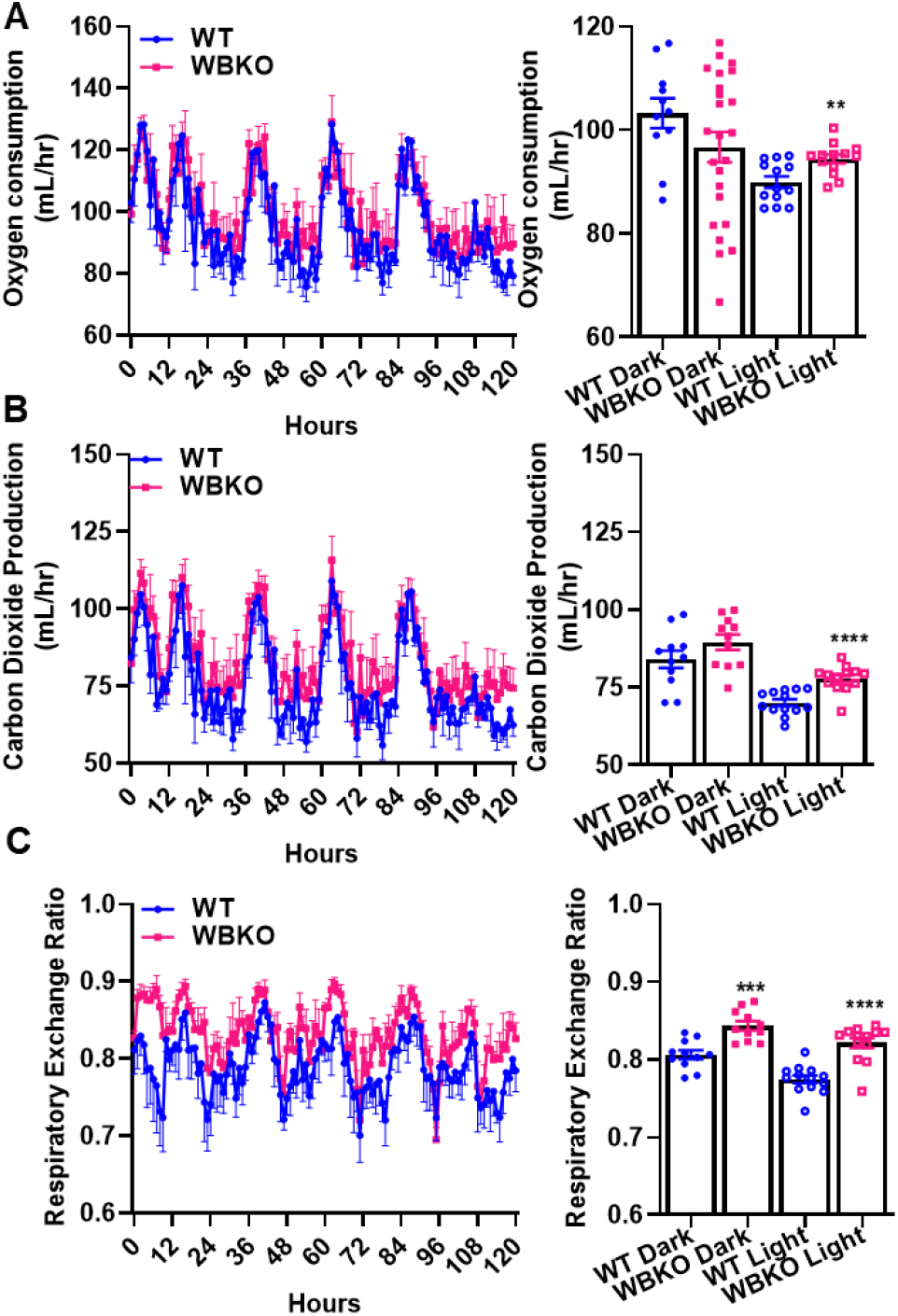
Altered energy metabolism in the WB-KO mice. Following 2-3 days of acclimation, metabolic parameters of the mice were measured by the Comprehensive Laboratory Animal Monitoring System (CLAMS). Measurements of VO2 (A) and VCO2 (B) gas exchanges were collected for five days. C) The respiratory exchange rate (RER) was obtained based on these values. Values are means ± SE; n= 4 (both female and male mice were included). Student t-test.

## Discussion

In this study, we showed that metal transporter Slc39a14/Zip14 regulates intestinal Zn homeostasis and gut microbiome composition, resulting in altered host metabolism. Deletion of the host gene encoding metal transporter, *Slc39a14/Zip14*, generated Zn-deficient epithelial and luminal content in the entire intestinal tract; reduced microbial diversity; altered host metabolome; and; shifted host energy metabolism toward glucose utilization (Fig. 7). To our knowledge, this is the first investigation that links the function of a mammalian metal transporter to gut microbiome composition, host serum metabolites, and energy metabolism.

**Figure 7.**
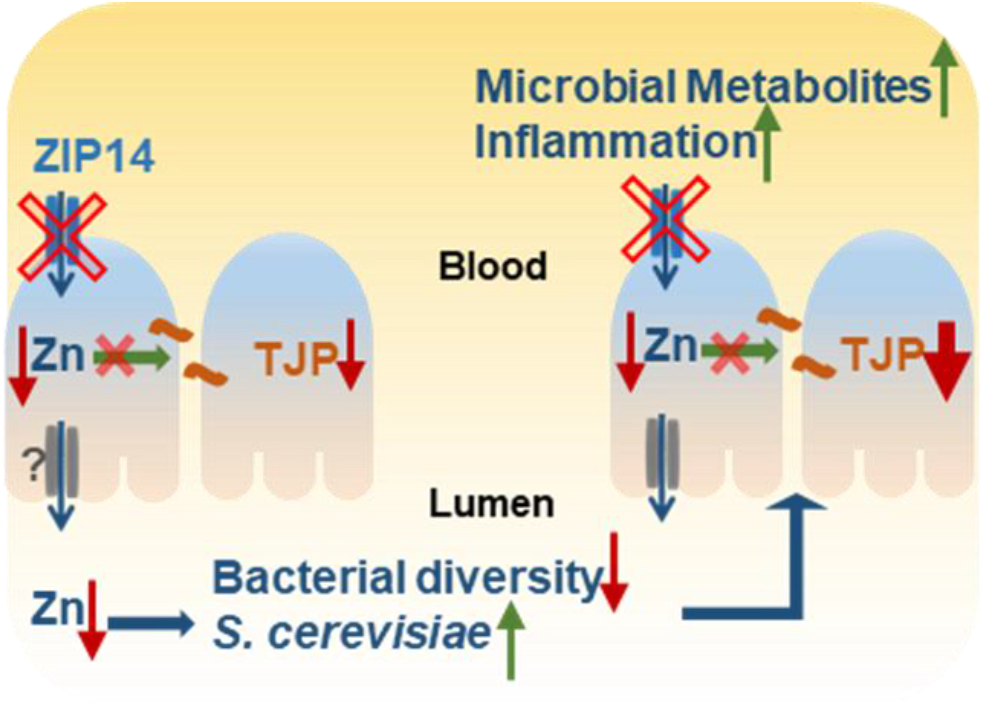
Model for a mechanism of intestinal permeability in *Zip14* KO mice. Deletion of the host gene encoding metal transporter, *Slc39a14/Zip14*, generates Zn-deficient epithelial and luminal content in the intestine. Reduced Zn concentrations in the enterocytes were shown to dysregulate tight junction proteins (TJP), subsequently increasing intestinal permeability ^68^. Reduced Zn concentrations in the lumen lower bacterial diversity and give the opportunity to *S. cerevisiae* overgrowth, further contributing to intestinal permeability. Consequently, altered host and microbial metabolites, inflammation, and shift in energy metabolism are found in the host.

Our previous studies showed localization of ZIP14 both to the basolateral side of the enterocyte and endosomes ^6^. In the *Zip14* KO mice, increased endosomal Zn concentrations suggested that Zn was entrapped in these compartments and generated Zn-deficient enterocytes. Here we showed that deletion of *Zip14* caused a deficiency in the intestinal tissue and the luminal content (Fig. 1A, B). These mice consumed a standard chow diet, and there was no difference in food consumption (data not shown). Therefore, low intestinal and luminal Zn suggested that Zn transport in serosal to mucosal direction was impaired in addition to endosomal entrapment in the absence of basolaterally localized ZIP14. ZIP4 is an apically localized Zn transporter and is responsible for the absorption of Zn. It is known to be upregulated during Zn deficiency to increase Zn absorption ^33^. ZnT1 on the basolateral side of the intestine was proposed to facilitate Zn transport from enterocytes to blood ^8^. ZnT1 is also regulated by Zn status ^5^. It was shown that during Zn deficiency, ZnT1 was downregulated to retain Zn in the cells. Upregulation of ZIP4 and downregulation of ZnT1 in isolated enterocytes from *Zip*14 KO mice further supported the Zn-deficient environment.

ZIP14, besides Zn, can transport manganese and iron ^21^. We have previously shown that Zn and manganese homeostasis were altered in *Zip14* KO mice at a steady-state ^18,20,23,34^, while ZIP14-related alterations in iron transport were largely observed in iron overload ^35^. Our current (Fig. 1) and previous results confirmed that both Zn and manganese transport in serosal to mucosal direction was impaired when *Zip14* was deleted ^23,34^. Since Zn, iron, and manganese are essential for microbial growth, the changes that we observe in gut microbiome composition could be caused by either Zn or Mn deficient intestine of *Zip14* KO mice. Manganese deficiency in humans is extremely rare ^36,37^. However, a rare genetic mutation in metal transporter, *ZIP8*, causes manganese deficiency with enhanced susceptibility to dextran sulfate sodium-induced injury ^15^. Contrary to manganese, Zn deficiency is recognized by WHO as a major global risk factor for disease mortality and morbidity ^38^. Importantly, Zn deficiency induces diarrhea and increases the risk of gastrointestinal diseases ^39,40^. Zn deficiency is found in IBD patients, and Zn supplementation studies have shown improved disease outcomes ^10^. However, human studies linking Zn, Zn transporters, gut microbiome composition, and intestinal health are lacking. In animal models, both Zn excess and deficiency were shown to modulate the microbiome profile. High dietary Zn in mice reduced the diversity of the gut microbiome community^41^, while in piglets, it caused either no change or increased microbiota diversity of ileal digesta and decreased microbiota diversity of the colonic digesta based on the dose and duration of Zn supplementation or the growth phase of the animals ^42^. A less diverse cecal microbiome composition was detected in gallus gallus in response to chronic dietary Zn deficiency ^43^, supporting our finding of lower bacterial α-diversity in the Zn-deficient intestine of *Zip14* KO mice.

The gut microbiome consists of bacteria, viruses, protozoa, and fungi in the gastrointestinal tract. The gut microbiota produces a broad range of metabolic products that impact host biology by acting on cells within the gastrointestinal tract or entering circulation and affecting distal sites within the body ^44,45^. Among the upregulated metabolites in the serum of *Zip14* KO mice (Fig. 3), N-acetyl-D-tryptophan, serotonin, and creatinine levels were shown to be different between germ - free and conventional mice ^46^, suggesting microbial origin/contribution. Enteric bacteria or yeast can generate tryptophan metabolites such as serotonin and N-acetyl-tryptophan ^47,48^. While either host or microbiome could make serotonin, N-acetyl-D-tryptophan could be unique to *Saccharomyces cerevisiae* (*S. Cerevisiae*) in the gut due to the production of a unique enzyme acting on D-amino acids, D-amino acid N-acetyltransferase ^29,30^. Lower bacterial diversity in the gut of *Zip14* KO mice could provide an opportunity for *S. cerevisiae* overgrowth. Of note, yeast cells survive during Zn deficiency due to multiple homeostatic and adaptive strategies ^49^. Increased intestinal permeability is observed both in Zn deficiency and inflammatory bowel disease (IBD) ^10^. Furthermore, Zn deficiency ^50^ and *S. cerevisiae* overgrowth ^51^ have been shown in IBD. Interestingly, serotonin was also shown to be upregulated in Crohn’s disease ^52^. These are, collectively, in agreement with the phenotype of *Zip14* KO mice with intestinal permeability and low-grade chronic inflammation.

Here we found a shift in energy metabolism towards continuous glucose utilization as an energy source with significantly greater TCA cycle intermediate metabolites in the serum of *Zip14* KO mice. We have previously shown that in the absence of *Zip14*, hepatic glucose uptake was increased along with greater glycogen stores ^53^ suggesting a greater amount of glucose available for energy production. The liver may have two ways of responding to this: increasing inefficient metabolism (Fig. 5-increased TCA cycle metabolites) and exporting these calories in the form of fat for deposition in peripheral tissues ^20^. Energy metabolism is also affected by inflammation. During chronic inflammation, adaptive immune cells, specifically T lymphocytes, go through metabolic reprogramming in favor of oxidative phenotype ^54^, in agreement with low-grade chronic inflammation and increased TCA cycle intermediates in *Zip14* KO mice. We have previously shown mitochondrial localization of ZIP14 in hepatocytes ^18^. In addition to metabolic reprogramming in adaptive immune cells of *Zip14* KO mice, the shift in energy metabolism could also be due to ZIP14-mediated regulation of mitochondrial metal concentration and the function of corresponding metalloenzymes. It is important to acknowledge that further studies with tissue/cell type-specific conditional *Zip14* KO mice along with dietary interventions for Zn and manganese are warranted to test these possible roles for ZIP14.

## Methods

### Animals and treatments

*Animals*. Heterozygous (*Zip14*^*+/-*^) mice are a generous gift of Dr. Robert J. Cousins. Heterozygous *Zip14*^*+/-*^ on the mixed background of C57BL6 and129S5 mice were bred to obtain homozygous whole-body (WB) *Zip14* KO (*Zip14*^*-/-*^) and WT (*Zip14*^*+/+*^) mice ^55^. The mice were maintained using standard rodent husbandry and received a commercial, irradiated chow diet (Harlan Teklad 7912; Indianapolis, IN, with 60 mg Zn/kg as ZnO) and tap water. Age-matched mice from both sexes were used as young adults (8–16 wk of age). Euthanasia was through exsanguination by cardiac puncture under isoflurane anesthesia. Protocols were approved by the Cornell University Institutional Animal Care and Use Committees. *Metabolic phenotyping*. Metabolic parameters of the mice were measured by the Comprehensive Laboratory Animal Monitoring System (CLAMS) by using Promethion (Sable Systems). Mice were transferred to the CLAMS metabolic cages and allowed to acclimate for 2-3 days with free access to food and water. Following the acclimation, measurements of VO_2_ and VCO_2_ gas exchanges and food and water consumption were collected for five days. The data were analyzed using Expedata software system (v1.9.27), Macro interpreter (v2.40), and then CalR software (v.1.2). ***Body composition***. Measurements were conducted in awake animals by time-domain nuclear magnetic resonance using the Minispec LF65 Body Composition Mice Analyzer. ***Fasting blood glucose***. Following overnight fasting, blood glucose levels were measured by tail bleeding using the OneTouch® Ultra Blood glucose monitoring system. ***Antibiotics treatment***. Antibiotics (Abx) (Ampicillin 1 g/L, neomycin 1 g/L, gentamycin 500 mg/L; penicillin 100U/L, vancomycin hydrochloride 1g/L, metronidazole 500mg/L) were given in the drinking water, ad libitum as described previously ^56,57^ for three weeks. Abx water was replaced with control water 4 hours before fecal transplant. ***Anti-fungal treatment***. Drinking water was supplemented with fluconazole (0.5 mg/mL, Sigma) for ten days ^58,59^. ***Fecal transplant***. Fecal pellets were collected from co-caged WT and WB KO mice to prepare the fecal slurry. 0.2 g of the fecal pellet was homogenized in 1mL of phosphate-buffered saline (PBS) using bullet blender (Next Advance). Following centrifuging at 800 x g for 3 min and filtering by 30 μm cell strainer, antibiotics-treated mice were inoculated for three consecutive days with 0.2 ml of the fecal slurry ^57,60^. Following conventionalization, metabolic phenotypes were assessed weekly by measuring fasting blood glucose and body composition by NMR. At the end of the three weeks, blood and tissue were collected for further biochemical analysis. ***Permeability Assay***. Following morning fasting (removal of food and water for 4 hours), mice were be given FITC-dextran (60 mg/100 g FITC-dextran 4) (Sigma-MW 3000-5000) via oral gavage. Blood was collected into EDTA tubes by cardiac puncture to obtain plasma at two hours post-gavage. FITC-dextran levels in plasma were measured fluorometrically at 485 nm emission/528 nm excitation.

### Analytical methods

#### Metal Assays

The entire excised small intestine was cut into two sections as proximal (the first 11cm) and distal (rest) and perfused with a metal chelating buffer (10 mM EDTA, 10 mM HEPES, and 0.9% NaCl_2_) (equal volume/section). Similarly, the colon was excised and perfused with the above buffer. The contents of the intestine and colon were centrifuged at 3000xg to obtain the lumen (supernatant) and digesta (pellet). Aliquots of tissue, lumen, digesta, and the fecal pellets were digested at 95°C overnight in HNO_3_ to measure metal concentrations using Microwave Plasma-Atomic Emission Spectrometry (MP-AES). Normalization was to tissue weight or volume (intestinal contents).

### Gut microbiome and serum metabolite profiling

Fresh fecal pellets were collected from mice that were co-caged for one month prior. ***DNA extraction and sequencing***. DNA samples were extracted from the mice fecal samples using DNeasy PowerSoil Kit (QIAGEN, Inc.) according to the manufacturer’s instruction. Region V4 of the 16S rRNA gene was amplified using the universal primers 515F and GoLay-barcoded 806R and the PCR program as previously described^61^. Samples were amplified in duplicate with the following thermocycler protocol: hold at 94°C for 3 min; 30 cycles of 94°C for 45 s, 50°C for 1 min, 72°C for 1.5 min; and hold at 72°C for 10 min, and the duplicate final amplified products were pooled. Amplicons were further purified using Mag-Bind® RxnPure Plus (Omega Bio-tek, Inc.) and quantified with NanoDrop 2000 spectrophotometer (Thermo Fisher Scientific), and 150 ng of amplicons from each sample were pooled and paired-end sequenced (2 × 250bp) on an Illumina MiSeq instrument. ***16S rRNA gene sequencing analysis***. Reads were demultiplexed using QIIME v1.9.1 (split_libraries_fastq.py) before denoising and processing with Divisive Amplicon Denoising Algorithm 2 (DADA2) ^62^. Taxonomy was assigned by mapping with the Greengenes 16S rRNA Gene Database ^63^. The QIIME output data was then imported to RStudio (Version 1.0.136) with the Bioconductor package phyloseq ^64^ to normalize and plot the input data. All statistical analyses were done in R using the vegan ^65^ and DESeq2 ^66^. A Benjamini–Hochberg FDR of 0.01 was used as the cutoff for statistical significance for heatmap plotting unless otherwise noted. ***Saccharomyces cerevisiae detection***. For isolating fungal DNA, an equal amount of colon digesta from each mouse were suspended in 50 mM Tris buffer (pH 7.5) supplemented with one mM EDTA, 0.2% b-mercaptoethanol and 1000 units/ml of lyticase (Sigma), incubated at 37°C for 30 min to disrupt fungal cells as described ^58,59^ prior to processing through the QIAamp Fast DNA Stool Mini Kit (Qiagen). The quantitative PCR (qPCR) reactions with an equal amount of DNA were set up using power track SYBR green master mix and run in Roche Lightcycler 480-II RT-PCR system. The S. cerevisiae-specific primer sequences were forward 5-AGGAGTGCGGTTCTTTG-3 and reverse 5-TACTTACCGAGGCAAGCTACA-3 ^67^.

#### Serum untargeted metabolomics profiling

Gas chromatography (GC) coupled to time-of-flight mass spectrometry (TOF-MS) was performed using Gerstel CIS4 –with dual MPS Injector/ Agilent 6890 GC-Pegasus III TOF MS by UC Davis West Coast Metabolomics Center. Statistical and pathway analyses were performed using MetaboAnalyst 5.0 ^31^. Data were normalized to the median to account for variance between samples and generalized logarithm transformed to account for differences in metabolite abundance. Metabolites with a fold change >1.5 and p < 0.05 were considered to be significantly different for pathway analysis. Pathway analysis was conducted using KEGG human metabolic pathways. Disease signature analysis was conducted using metabolite sets reported in human blood.

### Isolation of Enterocytes and Western blot analysis

Lumenal contents of the proximal intestine were flashed by PBS prior to evert them using bamboo sticks with pointed ends. Everted intestines were incubated in PBS with 1.5 mM EDTA solution for 20 min and released enterocytes were collected by centrifugation at 4°C at 500 x g for 5 min. Following two washes, the pellets were resuspended in lysis buffer (Tris-HCl, 137mM NaCl, 10% glycerol, 1% Triton X-100, 2 mM EDTA) with protease and phosphatase inhibitors (Thermo Scientific) added along with the PMSF (Sigma-Aldrich). During enterocyte separation, all the buffers were supplemented with protease inhibitors (AGScientific). Solubilized proteins were separated by 10% SDS-PAGE. Visualization was done by chemiluminescence (SuperSignal, Thermo Fisher) and digital imaging (Protein Simple). Rabbit anti-mouse ZIP14 antibody was custom made by Genscript. Antibodies for ZIP4, ZnT1, and GAPDH were obtained from Proteintech (20625), Novusbio (BP-94196), and Cell Signaling (14C10), respectively.

## Statistical Analyses

Data are presented as means ± SE. Significance was assessed by Student’s t-test for single comparisons. Statistical significance was set at P < 0.05. Analyses were performed using GraphPad Prism. The statistical criteria for metabolomics analysis and 16S rRNA gene sequencing analysis were included in the methods.

